# A quinolin-8-ol sub-millimolar inhibitor of UGGT, the ER glycoprotein folding quality control checkpoint

**DOI:** 10.1101/2022.06.21.496940

**Authors:** Kevin P. Guay, Roberta Ibba, JL Kiappes, Maria De Benedictis, Ilaria Zeni, James D. Le Cornu, Mario Hensen, Anu V. Chandran, Anastassia L. Kantsadi, Alessandro T. Caputo, Juan I. Blanco Capurro, Yusupha Bayo, Johan C. Hill, Kieran Hudson, Andrea Lia, Snežana Vasiljević, Carlos P. Modenutti, Stephen G. Withers, Marcelo Martí, Emiliano Biasini, Angelo Santino, Daniel N. Hebert, Nicole Zitzmann, Pietro Roversi

## Abstract

The Endoplasmic Reticulum (ER) glycoprotein folding Quality Control (ERQC) machinery aids folding of glycoproteins in the ER. Misfolded glycoprotein recognition and ER-retention is mediated by the ERQC checkpoint enzyme, the 170 kDa UDP-Glucose glycoprotein glucosyltransferase (UGGT). UGGT modulation is a promising strategy for broad-spectrum antivirals, rescue-of-secretion therapy in rare disease caused by responsive mutations in glycoprotein genes, and many cancers, but to date no selective UGGT inhibitors are known. Towards the generation of selective UGGT inhibitors, we determined the crystal structures of the catalytic domain of *Chaetomium thermophilum* UGGT (*Ct*UGGT_GT24_), alone and in complex with the inhibitor UDP-2-deoxy-2-fluoro-D-glucose (U2F). Using the *Ct*UGGT_GT24_ crystals, we carried out a fragment-based lead discovery screen via X-ray crystallography and discovered that the small molecule 5-[(morpholin-4-yl)methyl]quinolin-8-ol (5M-8OH-Q) binds a *Ct*UGGT_GT24_ ‘WY’ conserved surface motif that is not present in other GT24 family glycosyltransferases. The 5M-8OH-Q molecule has a 613 *µ*M binding affinity for human UGGT1*in vitro* as measured by saturation transfer difference NMR spectroscopy. The 5M-8OH-Q molecule inhibits both human UGGT1and UGGT2 activity at concentrations higher than 750 *µ*M in modified HEK293-6E cells. The compound is toxic *in cellula* and *in planta* at concentrations higher than 1 mM. A few off-target effects are also observed upon 5M-8OH-Q treatment. Based on an *in silico* model of the interaction between UGGT and its substrate *N* -glycan, the 5M-8OH-Q molecule likely works as a competitive inhibitor, binding to the site of recognition of the first GlcNAc residue of the substrate *N* -glycan.

**Significance Statement:** When a candidate drug target is the product of a housekeeping gene - i.e. it is important for the normal functioning of the healthy cell – availability of inhibitors for tests and assays is of paramount importance. One such housekeeping protein is UGGT, the enzyme that makes sure that only correctly folded glycoproteins can leave the endoplasmic reticulum for further trafficking through the secretory pathway. UGGT is a potential drug target against viruses, in certain instances of congenital rare disease, and against some cancers, but no UGGT inhibitors are known yet. We discovered and describe here a small molecule that binds human UGGT1 *in vitro* and inhibits both isoforms of human UGGT *in cellula*. The compound paves the way to testing of UGGT inhibition as a potential pharmacological strategy in a number of medical contexts.

---

**I**n the endoplasmic reticulum of eukaryotic cells, the ER glycoprotein folding Quality Control (ERQC) system ensures ER retention of immature glycoproteins and assists their folding (1). Glycoprotein ERQC is central to glycoproteostasis, which in turn plays a major role in health and disease (2, 3). Glycoprotein ERQC is reliant on detection of glycoprotein misfolding, effected by its checkpoint enzyme, UDP-glucose glycoprotein glucosyltransferase (UGGT). UGGT is capable of detecting non-native and slightly misfolded glycoproteins, and re-glucosylates its clients to flag themNfor ER retention(4).

While other components of ERQC have been studied as drug targets(5–7), cellular consequences of pharmacological modulation of UGGT have been relatively understudied - partly because of the risks associated with targeting core cell housekeeping machineries, and partly because there are no known UGGT selective inhibitors. UGGT is inhibited by its product UDP(8) and by the non-hydrolysable UDP-Glucose (UDP-Glc) analogue UDP-2-deoxy-2-fluoro-D-glucose (U2F) but neither molecule is specific. Selective and potent UGGT modulators would be important reagents for cell biology for interrogating the cell biology of the secretory pathway, as well as having therapeutic potential in several areas of medical science, such as virology(9–11), rare genetic disease (12, 13) and cancer(14–16). Selective and potent UGGT activity modulators also have potential applications in biotechnology and agricultural science (17–20).

We set out to search for ligands of UGGT by fragment-based lead discovery (FBLD) using X-ray crystallography(21–24). No crystal structures of mammalian UGGTs have been obtained so far, but atomic resolution structures of UGGTs from thermophilic fungi have been determined(25–27). Although compounds binding the N-terminal, misfold-recognising portion of the enzyme would also be potential UGGT inhibitors, we decided to target the C-terminal catalytic domain, given the high 70% similarity and 60% identity between human and fungal sequences in this portion of the enzyme. In our FBLD effort, each of hundreds of crystals of the catalytic domain of *Chaetomium thermophilum* UGGT (*Ct*UGGT) was soaked with a different chemical compound from a molecular fragment library.

The study yielded the first small molecule UGGT ligand, 5-[(morpholin-4-yl)methyl]quinolin-8-ol (5M-8OH-Q for short in what follows), with sub-millimolar affinity for human UGGT1. The 5M-8OH-Q molecule inhibits both human paralogues of UGGT, UGGT1and UGGT2, at concentrations higher than 750 *µ*M in modified HEK293-6E cells. A medicinal chemistry program to generate more potent and selective UGGT inhibitors starting from 5M-8OH-Q is in progress.

## Ct UGGT_GT24_ and ^U2F^*Ct*UGGT_GT24_ crystal structures

FBLD by X-ray crystallography requires the growth of hundreds of well-diffracting crystals of the target macromolecule. None of the crystals of full-length UGGT we grew so far diffracted past 2.8 Å(25, 27), but 1.35 and 1.4 Å crystal structures of the catalytic domain of *Thermomyces dupontii* UGGT (*Td*UGGT), in complex with UDP and UDP-Glc, respectively, have been described(26). Inspired by those results, we cloned the catalytic domain of *Ct*UGGT without its C-terminal ER-retrieval motif (hereinafter *Ct*UGGT_GT24_) in the pHLsec vector for secreted mammalian expression(28). The domain belongs to the GT24 family of glycosyltransferases(29). *Ct*UGGT_GT24_ protein was then expressed, purified, and its structure determined by X-ray crystallography. A crystal structure of *Ct*UGGT_GT24_ with the inhibitor U2F was also determined. Tables S1 and S2 list the X-ray data collection statistics and structure refinement statistics, respectively.

Half of the coordination sphere of the Ca^2+^ ion in the *Ct*UGGT_GT24_ active site is common to both structures: the side chains of D1302 and D1304 (belonging to the UGGT conserved ^1302^DAD^1304^ motif) and the side chain of the conserved D1435 always take up three invariant coordination sites around the Ca^2+^ ion (Figures SI1-A,B). In the 1.8 Å structure of apo *Ct*UGGT_GT24_ (PDB ID 7ZKC) two water molecules occupy two of the three remaining coordination sites around the Ca^2+^ ion, with the main chain carbonyl oxygen of L1436 completing the ion’s octahedral coordination (Figure S1-A).

The *Ct*UGGT_GT24_ structure in complex with U2F (structure ^U2F^*Ct*UGGT_GT24_, PDB ID 7ZLU) captures a conformation of U2F likely equivalent to that following initial UDP-Glc binding to UGGT: the U2F ribose ring points towards the solvent (Figures S1-B and C). The uracyl ring O4 atom accepts a hydrogen bond from the main chain NH of S1207 and its N3 atom donates one hydrogen bond to the main chain O of the same residue (Figures S1-B and C). The U2F uracyl ring forms a *π*-stacking interaction with the conserved *Ct*UGGT Y1211 whose side chain rotates slightly when compared to the apo structure, to accommodate the ligand. Rearrangement of the Ca^2+^ ion coordination sphere with respect to the apo structure is also observed; the two apo waters are replaced by an O atom from the *β* phosphate and by the O2 atom of the Glc ring of U2F. The main chain of L1436 moves away from the Ca^2+^ ion and a water molecule occupies its Ca^2+^ coordination site (Figures S1-B and C). The U2F pose suggests that the UGGT active site selects UDP-Glc over UDP-Gal(30–32): in UDP-Glc the glucose O4 atom forms hydrogen bonds to the side chains of conserved W1280 and D1396, but these interactions would be lost in UDP-Gal, because of the difference in stereochemistry between Glc and Gal in position 4 (see Figure S1-C).

## 5M-8OH-Q is a novel *Ct*UGGT_GT24_ ligand

The *Ct*UGGT_GT24_ crystals (grown with UDP-Glc and Ca^2+^) were used for soaking hundreds of compounds to find *Ct*UGGT_GT24_ ligands by FBLD. The best hit was 5-[(morpholin-4-yl)methyl]quinolin-8-ol (5M-8OH-Q), a crystal soaked with which diffracted to 2.5 Å. To confirm binding, we then grew a co-crystal of *Ct*UGGT_GT24_ with the 5M-8OH-Q molecule and obtained a 1.7 Å structure (^5M-8OH-Q^*Ct*UGGT_GT24_, PDB ID 7ZLL). The compound binds to a conserved patch on the surface of the *Ct*UGGT_GT24_ domain, about 15 Å away from the UDP-Glc binding site (Figure 1A and Figure S2-A). The morpholine ring is partially disordered in the crystal, but one of its ring placements is Å from the conserved ^1396^DQD^1398^ motif coordinating the Glc ring of UDP-Glc (Figure 1A); the ligand also causes a displacement of the side chain of *Ct*UGGT_GT24_ ^1346^Y. Through this displacement, the 8OH-quinoline ring inserts and is sandwiched between the aromatic side chains of the conserved residues ^1346^YW^1347^ - which we propose to call the ‘YW clamp’. The two aromatic side chains stabilize the quinoline ring forming an aromatic trimer (33); the 8-OH group of the quinoline also establishes an H-bond to the side chain of ^1402^H and a *π*-interaction is visible between the quinoline ring N atom and the side chain of ^1350^Y (Figure 1A and Figure S2-B).

**Fig. 1.**
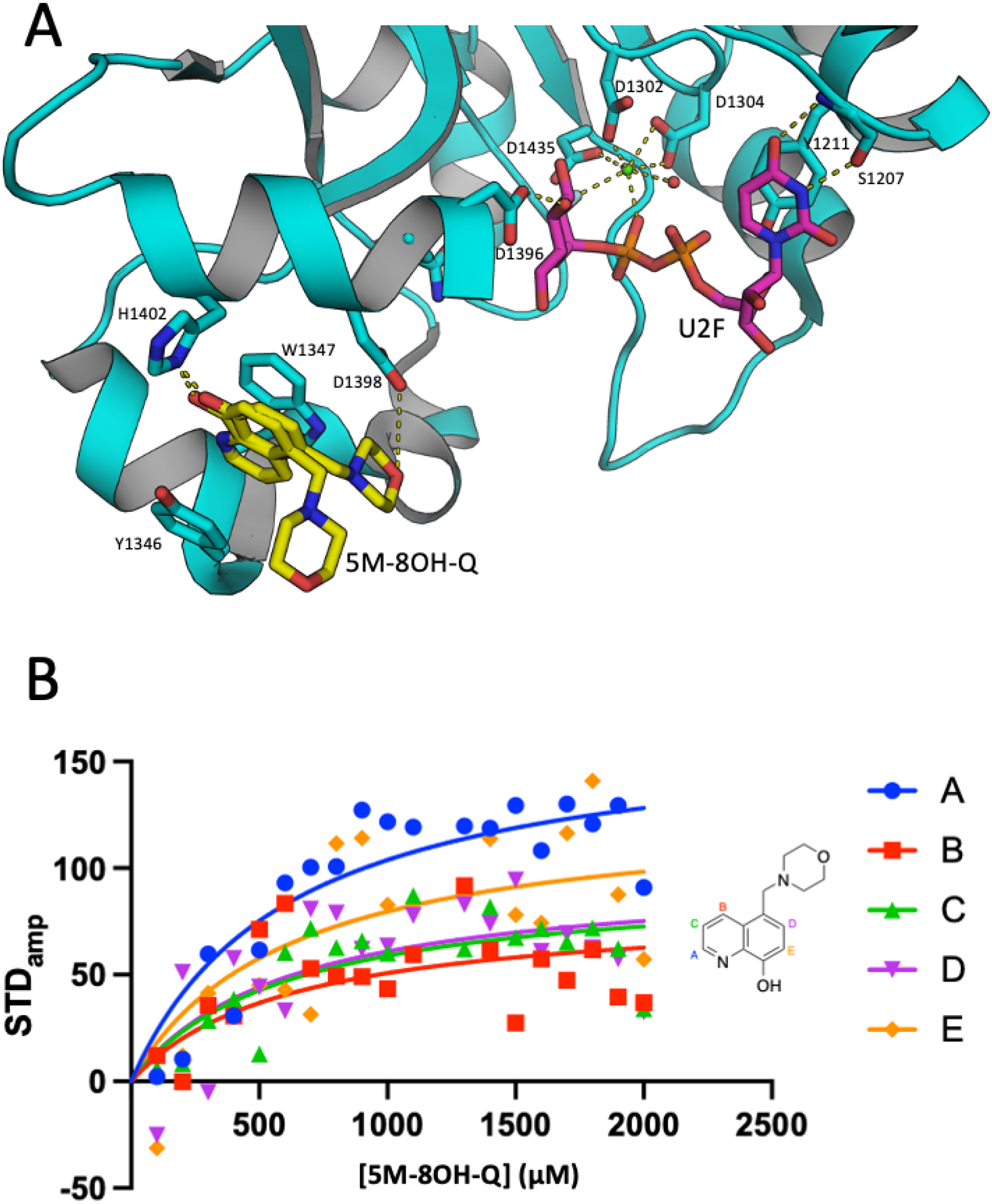
The active site of UGGT and the 5M-8OH-Q inhibitor. **A**: The active site of the ^U2F^*Ct*UGGT_GT24_ structure with the 5M-8OH-Q ligand from the overlayed ^5M-8OH-Q^*Ct*UGGT_GT24_ structure. Protein atoms in sticks representation; C cyan (but UDP-Glc(UDP) C magenta and 5M-8OH-Q C atoms yellow), O red, N blue, P orange, F light green. H-bonds and Ca^2+^-coordination bonds are in yellow dashed lines. The Ca^2+^ ion is a green sphere and its coordinating water molecules are red spheres. The side chains of residues D1302, D1304 and D1435 coordinate the Ca^2+^. The *Ct*UGGT ^1346^ YW^1347^ clamp, the conserved ^1346^ DQD^1347^ motif, H1402, Y1211 and the main chain of S1207 are in stick representation. Two of the poses of the 5M-8OH-Q inhibitor are shown. **B**: Measurement of the *K*_*d*_ dissociation constant of the complex between 5M-8OH-Q and human UGGT1 by STD NMR *in vitro*. Each curve follows the interaction of one of the H atoms of 5M-8OH-Q with the human UGGT1 protein.

## 5M-8OH-Q has sub-millimolar affinity for human UGGT1 *in vitro*

Saturation transfer difference (STD) NMR spectroscopy was used to measure the affinity of 5M-8OH-Q for human UGGT1*in vitro*. The concentration dependence of the STD amplification factor (STD_amp_) was measured on a 500 MHz NMR spectrometer for each of the 5 aromatic hydrogen atoms in 5M-8OH-Q and the human UGGT1: 5M-8OH-Q K_d_ estimated for each of the datasets (Figure 1B). The fit to the five sets of data pertaining to each of the 5M-8OH-Q H nuclei was compatible with a single value of K_d_ of 613 *µ*M. Epitope mapping of the STD data is in good agreement with the crystal structure, with protons in the A and E environments having the largest STD_amp_ values, indicating they are in closest proximity to the protein when bound (Figure 1B).

## 5M-8OH-Q is a sub-millimolar inhibitor of human UGGTs *in cellula*

To ascertain if 5M-8OH-Q can be delivered to the ER and inhibit UGGT-mediated glucosylation *in cellula*, modified modified HEK293-6E cells were treated with the inhibitor, monoglucosylated glycoproteins isolated by GST-calreticulin (GST-CRT) affinity precipitation and the eluate analyzed by immunoblotting (34, 35). To ensure the CRT interaction resulted from UGGT glucosylation, and not from the initial glycan trimming that occurs during normal glycan maturation, CRISPR/Cas9 was used to knock out the *ALG6* gene: in *ALG6*^-/-^cells, the CRT lectin pulldown selects only monoglucosylated glycoproteins that were glucosylated by UGGT and not the ones produced by the ER glucosidases initial glycan trimming - as in this background the *N* -glycan precursors added to nascent glycoproteins initially lack the three glucoses(36) ^∗^.

In order to decide on the maximum assay concentration of 5M-8OH-Q, toxicity assays were carried out. In a trypan blue assay, toxic effects were observed around 1-2 mM 5M-8OH-Q and above in modified HEK293-6E cells: after 5 hours of treatment with 1 or 2 mM 5M-8OH-Q the viability was about 75-80% (Figure S4). This level of toxicity is comparable to the one observed upon 5M-8OH-Q treatment of *Arabidopsis thaliana (At)* plants, which are a good model for the study of ER glycoprotein folding quality control (38, 39) (Figure S5).

The *ALG6*^-/-^ HEK293-6E cells were treated with increasing concentrations of 5M-8OH-QandN-following incubation with the molecule - glucosylation of known UGGT substrate glycoproteins was analyzed by isolating monoglucosylated glycoproteins from the cell lysate. After GST-CRT precipitation, the eluate was probed for two known substrates of UGGT: the proprotein of human IFGR1 (ProIFGR1, a UGGT1substrate(35)) and the proprotein of HexB (ProHexB, a UGGT2 substrate(35)) and their glucosylation levels quantified. The amount quantified in each GST-CRT pulldown was divided by the total amount found within the sample’s whole cell lysate (WCL), resulting in the percent glucosylation at that dose of 5M-8OH-Q(35).

Levels of monoglucosylated IGF1R and HexB in the *ALG6*^-/-^ HEK293-6E cells decrease as the concentration of 5M-8OH-Q increases (Figure 2A, even-numbered lanes 2-18). In particular, a significant decrease in IGF1R and HexB glucosylation is observed at 0.5 and 0.75 mM 5M-8OH-Q, respectively, in agreement with the proposed K_*d*_ of 0.613 mM of human UGGT1 for 5M-8OH-Q as determined by STD-NMR. IGF1R and HexB glucosylation decrease from ∼17% to ∼4% and ∼9% to ∼2%, respectively, going from no treatment to 2 mM 5M-8OH-Q (Figure 2B-D). Interestingly, the overall levels of IGF1R and HexB glycoproteins also seem to decrease with increasing levels of 5M-8OH-Q (WCL lanes in Figure 2A). This could be explained by UGGT inhibition causing misfolding and ER associated degradation (ERAD) of its client glycoproteins, thus lowering their levels; alternatively, the side effect may be due to lack of specificity of 5M-8OH-Q at these relatively high concentrations.

**Fig. 2.**
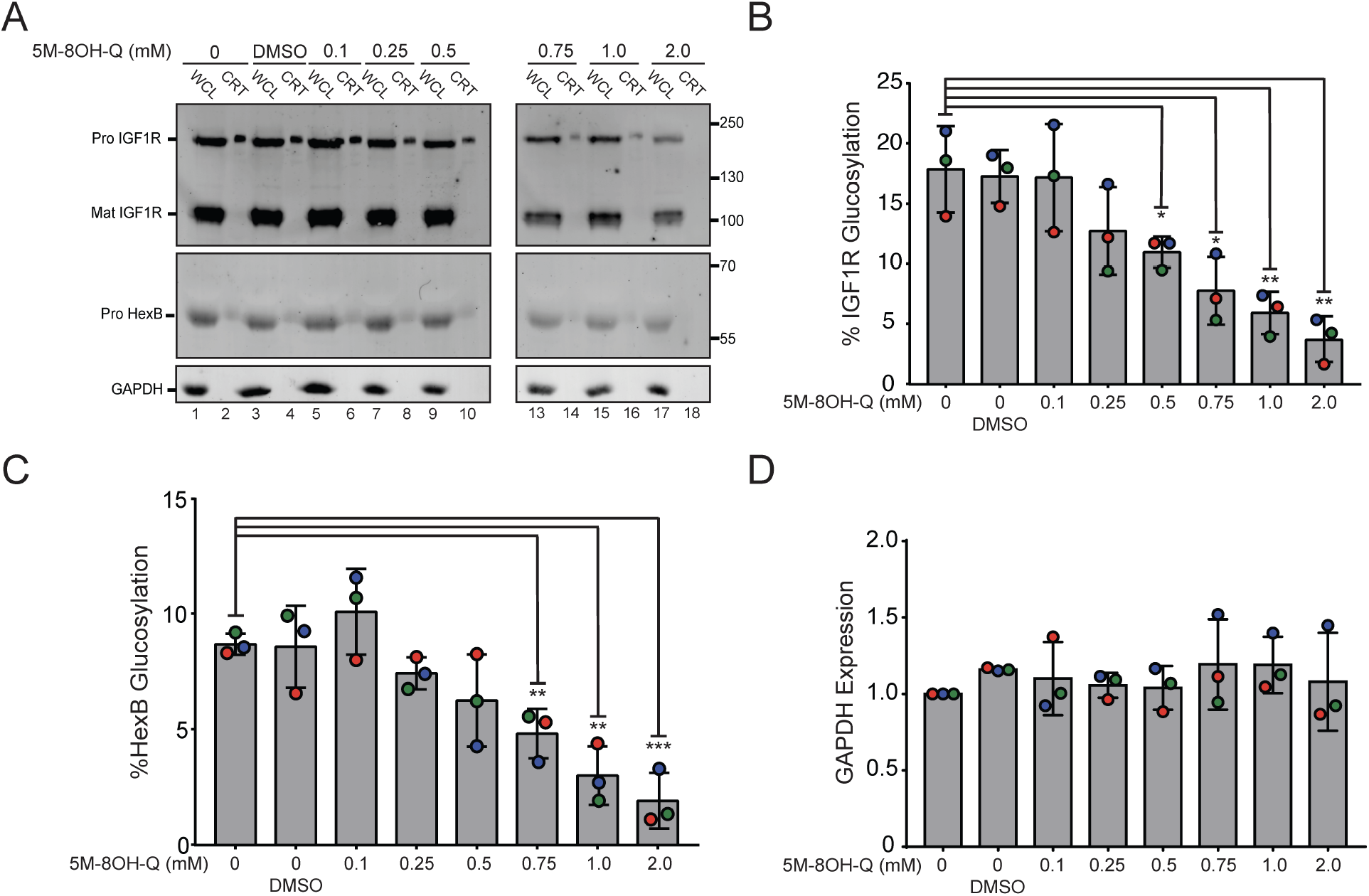
5M-8OH-Q dose-dependent inhibition of UGGT. **A**: *ALG6*^-/-^ HEK293-6E cells were cultured and treated with increasing concentrations of 5M-8OH-Q. The “0 mM” group was treated with no drug or vehicle. The vehicle control group was incubated with DMSO. The lysate was split between a whole cell lysate sample (20%, “WCL”) and a GST-CRT pulldown sample (60%, “CRT”), and resolved by 9% SDS-PAGE gel electrophoresis, before transferring the protein bands to a PVDF membrane. Imaged are immunoblots probed for IGF1R (whose proprotein *Hs*ProIFGR1 is a UGGT1 substrate(35)), HexB (whose proprotein *Hs*ProHexB is a UGGT2 substrate(35)) and GAPDH (loading control). Each data point comes from three independent biological replicates. **B**,**C**: Quantification of *Hs*ProIFGR1 and *Hs*ProHexB glucosylation over increasing amounts of 5M-8OH-Q from the experiments in **A**. Percent glucosylation was calculated by dividing the normalized CRT value by the normalized value from the WCL and multiplying by 100. **D**: Anti-GAPDH blot control. Protein samples were loaded to match the protein in the “0 mM” group for each condition. Error bars represent the standard deviation. Statistical significance levels: *: P *≤* 0.05; **: P *≤*0.01; ***: P *≤* 0.001.

Next, we asked whether 5M-8OH-Q inhibits both human paralogues of UGGT (UGGT1 and UGGT2 (35, 40, 41)). Double knock-out (KO) cells (*ALG6/UGGT1*^-/-^ and *ALG6/UGGT2*^-/-^ (35)) were exposed to 1 mM of the drug to measure glucosylation of IGF1R and HexB as described above (Figure 3A). As expected, glucosylation of IGF1R (a UGGT1 substrate) is significantly inhibited in both the *ALG6*^-/-^ and *ALG6/UGGT2*^-/-^, but not in the *ALG6/UGGT1*^-/-^ (Figure 3B). Similarly, glucosylation of the UGGT2 substrate HexB is inhibited in the *ALG6*^-/-^ and *ALG6/UGGT1*^-/-^ cells, but not in the *ALG6/UGGT2*^-/-^ cell line (Figure 3C). The levels of inhibition within each of these UGGT KO cell lines agree well with the findings described earlier (Figure 2 and (35)). In agreement to what is observed in Figure 2A, 5M-8OH-Q also decreases the levels of IGF1R and HexB in the WCL lanes (Figure 3A). Taken together these results suggest 5M-8OH-Q can reach the ER and inhibit both paralogues of UGGT.

**Fig. 3.**
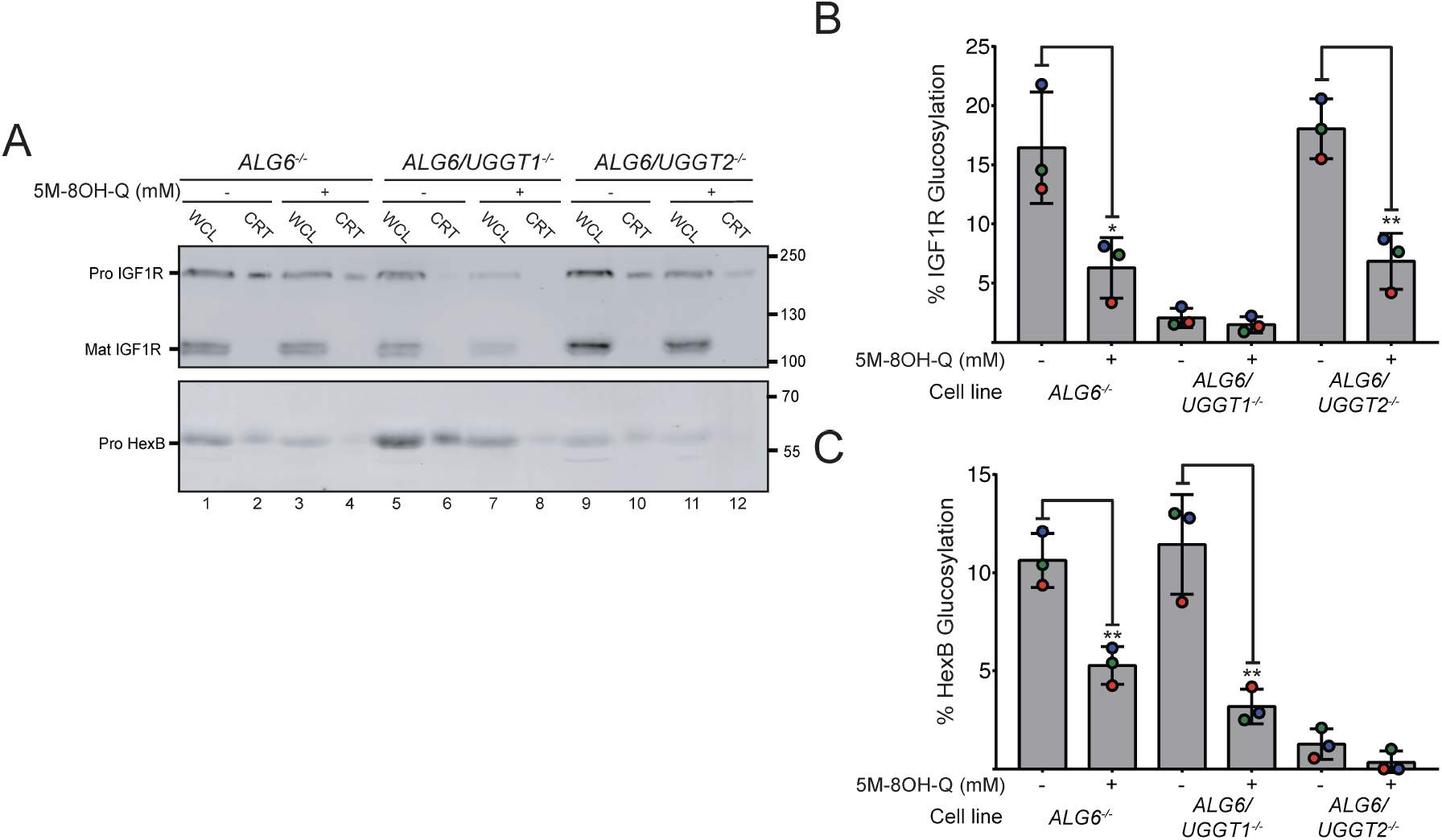
5M-8OH-Q inhibits both UGGT1 and UGGT2. **A**: *ALG6*^-/-^, *ALG6/UGGT1*^-/-^ and *ALG6/UGGT2*^-/-^ HEK293-6E cells were cultured and either not treated or treated with 1 mM 5M-8OH-Q to determine if the drug inhibits one or both of UGGT1 and UGGT2. After the cells were incubated with the inhibitor, they were lysed and split between a whole cell lysate sample (20%, “WCL”) and a GST-CRT pulldown sample (60%, “CRT”), and resolved by 9% SDS-PAGE gel electrophoresis, before transferring the protein bands to a PVDF membrane. Imaged are immunoblots probed for IGF1R (UGGT1 substrate) and HexB (UGGT2 substrate). Glucosylation of human ProIFGR1 and human ProHexB was observed in *ALG6*^-/-^ and *ALG6/UGGT2*^-/-^ and in the *ALG6*^-/-^ and *ALG6/UGGT1*^-/-^ cell lines, respectively. Each data point represents three independent biological replicates. **B**,**C** Quantification of human ProIFGR1 and human ProHexB glucosylation from **A**. Percent glucosylation was calculated by dividing the normalized value from the CRT lane by the normalized WCL. The resulting value was multiplied by 100 to obtain percent glucosylation. Error bars represent the standard deviation. Statistical significance levels: *: P ≤ 0.05; **: P ≤0.01; ***: P ≤ 0.001.

## *In silico* docking suggests that 5M-8OH-Q competes for the binding site of the first GlcNAc of the *N* -linked glycan

To gain insight into how 5M-8OH-Q inhibits UGGT, we built an *in silico* model of the Man_9_GlcNAc_2_ glycan bound to *Ct*UGGT using a combination of knowledge-based docking and Molecular Dynamics (see Material and Methods). For the positioning the acceptor Man ring of branch A of the glycan next to the UDP-Glc glucose ring (Figure 4A), we took advantage (see Material and Methods) of our U2F structure and of the UDP-Glc bound one (26).

**Fig. 4.**
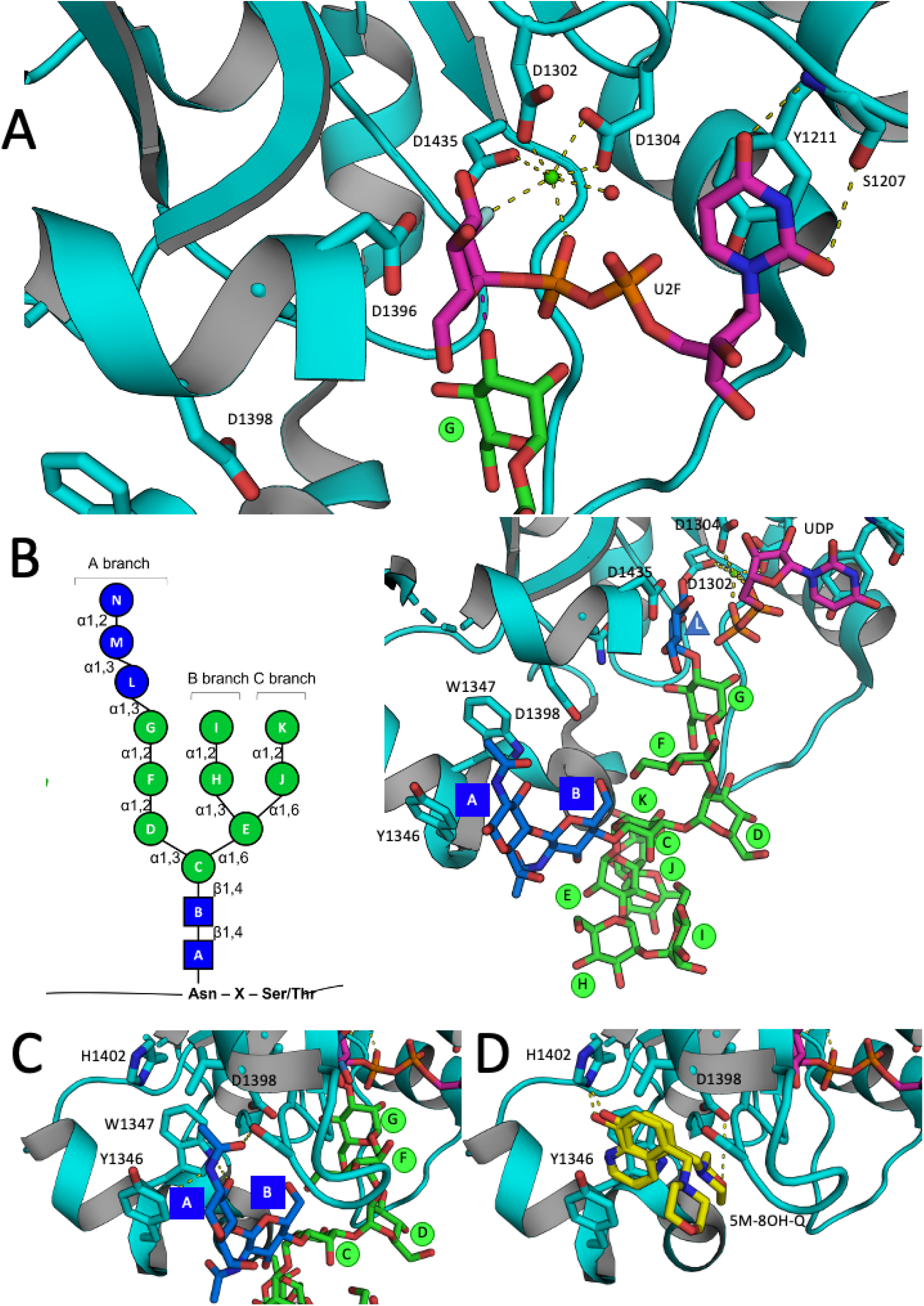
Modelling of the GlcNAc_2_Man_9_ glycan bound to *Ct*UGGT_GT24_ domain. **A**: Man “G” placement next to the UDP-Glc binding site, in an orientation suitable for the nucleophilic attack of its O3 oxygen to the glucose anomeric centre (red dashed line), to yield the *β*(1-3) Glc-Man bond. **B**: GlcNAc_2_ Man_9_ Glc_3_ glycan glycan nomenclature and final model of the GlcNAc_2_Man_9_Glc_1_ glycan docked onto the *Ct*UGGT_GT24_ domain. Saccharide moieties are colour-coded according to the scheme on the left hand side(61). **C**,**D**: The docked GlcNAc_2_ moiety of the N-linked glycan and 8-OH-Q share a binding pocket.

The surface of the UGGT catalytic domain on which the glycan docks according to our model is highly conserved across eukaryotic UGGT1s and UGGT2s(42). The A branch of the Man_9_GlcNAc_2_ glycan stretches towards the UGGT active site, while B and C branches point towards the solvent, fitting into shallower grooves, binding the protein with fewer interactions (Figure 4B). These observations are consistent with previous work showing that UGGT is able to glucosylate misfolded glycoproteins bearing GlcNAc_2_Man_8_ (Man “I” trimmed) and GlcNAc_2_Man_7_ (Man “I” and Man “K” trimmed) glycans (see Figure 4B)(43).

Importantly, the model suggests how UGGT recognises the first GlcNAc: the glycan’s first *N* -acetamide group faces directly into the hydrophobic cavity formed by residues Y1346, W1347, and L1392, its acetyl oxygen hydrogen-bonded to the L1392 backbone nitrogen and the S1391 hydroxyl group (Figure 4C), in agreement with the finding that the first GlcNAc is required for the Man_9_GlcNAc_2_ glycan to bind to UGGT (43, 44). As this is also the site of binding of 5M-8OH-Q (Figure 4D), the model suggests that the molecule acts as a UGGT competitive inhibitor with respect to the glycoprotein substrate.

## Discussion

Since its discovery in 1989(45), UGGT retains a central role in the standard model of glycoprotein ERQC. As such, and given the importance of glycoprotein folding to health and disease (3), UGGT is a potential target for drugs to treat a variety of conditions (12, 15, 46). As of today, no UGGT inhibitors have been described, apart from its product, UDP(47) - and the UDP-Glc analogue U2F - hardly good scaffolds for selective drug design, given that all genomes encode a plethora of proteins carrying a UDP- or a UDP-Glc-binding site. Until the molecular mechanisms underpinning misfold recognition are elucidated, and the portions of UGGT involved in this process are discovered(27), the catalytic domain remains the most promising target for novel classes of compounds that inhibit UGGT-mediated glucosylation of misfolded glycoproteins in the ER.

We grew crystals of *Ct*UGGT_GT24_ in order to hunt for novel ligands by FBLD. 8-OH-quinolines can chelate a great number of cations, including Cu^2+^, Bi^2+^, Mn^2+^, Mg^2+^, Fe^3+^, Al^3+^, Zn^2+^ and Ni^3+^(48), and synthesis of the 5M-8OH-Q ligand discovered through our FBLD effort was indeed originally described as part of studies of soluble aluminum complex dyes(49) or fluorescent Zinc sensors(50). In the medical field, 8-hydroxyquinoline derivatives can be used as insecticides, antibacterial, fungicidal, neuroprotective, and anti-HIV agents(51, 52). For example, the 5M-8OH-Q K_d_ for SARS-CoV-2 × main viral protease was estimated as 28.6 10^−6^ M by a recent *in silico* study (53).

Our ^5M-8OH-Q^*Ct*UGGT_GT24_ crystal structure shows that 5M-8OH-Q binds a conserved pocket on the surface of the protein, not far from the UDP-Glc binding site. *In vitro*, 5M-8OH-Q binds to full-length human UGGT1 with sub-millimolar K_d_. *In cellula* experiments show a concentration-dependent decrease in mono-glucosylation of the *Hs*ProIFGR1 and *Hs*ProHexB UGGT substrates upon treatment of HEK293-6E cells with 5M-8OH-Q (Figure 2), indicating that the molecule crosses the plasma and ER membranes and inhibits ER lumenal UGGTs.

Both UGGT isoforms are inhibited (Figure 3), a result that agrees with the conservation of the 5M-8OH-Q binding site in the catalytic domain of the two proteins. Besides *Hs*UGGT1 and *Hs*UGGT2, the human genome encodes 10 more genes containing GT-A and GT-B glycosyltransferase domains, but from sequence alignment, it appears that the YW clamp providing the 5M-8OH-Q binding platform is specific to UGGTs (GT24 family(54)), Figure S3). Our model of the Man_9_GlcNAc_2_ glycan bound to the catalytic domain of *Ct*UGGT shows the 5M-8OH-Q binding site partially overlapping with the Man_9_GlcNAc_2_ glycan binding site, which is also shared between UGGT1 and UGGT2(42).

The dose-dependent reduction of the levels of the UGGT substrates (*Hs*ProIFGR1 and *Hs*ProHexB)) observed in *ALG6*^-/-^ HEK293-6E cells (“WCL” lanes in Figure 2) could be due to indirect effects of UGGT inhibition: both client glycoproteins fold under UGGT control, so that UGGT inhibition may affect the levels of fully folded *Hs*ProIFGR1 and *Hs*ProHexB as well as UGGT-mediated glucosylation of the same clients. 8-OH quinolines are known to bind to a dozen mammalian proteins (see Table S3): for example, the quinoline N atom and the 8-OH substituent chelate metal ions and bind to catalytic sites of human demethylases, 2-oxoglutarate/iron dependent oxygenases, and *α*-ketoglutarate-dependent RNA demethylases (55–57). 5M-8OH-Q is toxic *in cellula* and *in planta* at concentrations higher than 1 mM (Figures S4 and S5). In preliminary assays of the effects of 5M-8OH-Q on secretion of other glycoproteins, we also observed altered levels of N-glycosylated proteins in the ER of *Arabidopsis thaliana* upon 2 mM 5M-8OH-Q treatment (Figure S6); and an increase of the alpha-cleavage of the human major prion protein (PrP) when treating HEK293 cells stably expressing it (Figure S7). At present it is unclear if the observed side effects are due to 5M-8OH-Q directly interacting with other proteins of the early secretory pathway, or to indirect effects of UGGT inhibition on UGGT glycoprotein clients’ folding and levels.

In summary, 5M-8OH-Q inhibits both UGGT1 and UGGT2 in cellula, targeting a site on the surface of the UGGT catalytic domain that is not present in other GT24 family glycosyltransferases. 5M-8OH-Q provides therefore a useful starting point for the synthesis of UGGT modulators for the treatment of diseases caused by “responsive mutants”, as persistent UGGT-mediated glucosylation may prevent trafficking of slightly misfolded, but otherwise functional, glycoproteins to their correct cellular locations(12). UGGT inhibition may one day also find application as an anti-cancer strategy, as some UGGT substrate glycoproteins (35) are selectively up-regulated in cancer cells(15). Replication of pathogenic enveloped viruses whose envelope glycoproteins fold under UGGT control may also be impaired by UGGT inhibitors(46). The strong conservation of UGGT sequence/function across eukaryotes (3) broadens the potential impact of such molecules to many fields: examples are plants as *in vivo* models to study secretion (38, 39, 58); stress-resistant genetically modified crops (59); or expression systems for recombinant glycoproteins (60). The relative low affinity and likely low specificity of 5M-8OH-Q are hardly surprising given that the molecule was discovered as a UGGT binding fragment during a FBLD effort and it has not been chemically modified to improve its potency and selectivity yet. A medicinal chemistry program that will yield and test the first generation of 5M-8OH-Q derivatives is in progress.

## Materials and Methods

### *Ct* UGGTprotein expression and purification

*Ct*UGGT was expressed and purified as described in (42).

### UGGT1 cloning, protein expression and purification

The C-terminally His-tagged construct encoding human UGGT1 residues 43-1551 was PCR-amplified from the commercially sourced vector UGGT1-pUC57 (GenScript) with primers: OPPF_UGGT1_Fwd: gcgtagctgaaaccggcGACTCAAAAGCCATTACAAC-CTCTCT OPPF_UGGT1_Rev: gtgatggtgatgtttTTTCTGAGGACCTTCTCGGCTTGG. These primers were designed to surround the insert with an N-terminal AgeI restriction site and a C-terminal KpnI site (after the C-terminal 6xHis tag and the stop codon). The amplified DNA was run on a 0.8% agarose gel and the correctly-sized fragment excised and purified using the QIAquick Gel Extraction Kit (QIAgen). The pOPINTTGneo plasmid was linearised with 20 units of both AgeI-HF and KpnI-HF restriction enzymes, incubated with 1x CutSmart Buffer (New England BioLabs) and 500ng of pHLSec DNA and digested at 37°C overnight. Both the linearised pOPINTTGneo and the UGGT1insert DNA were run on a 0.8% agarose gel and the correctly-sized fragments excised and purified using the QIAquick Gel Extraction Kit (QIAgen). DNA ligation of the linearised pOPINTTGneo vector and the human UGGT1insert was achieved by In-fusion™ligation-independent cloning (Takara Ltd.)

Transfection of HEK293F cells with the UGGT1-pOPINTTGneo plasmid and expression of the recombinant human UGGT1protein were carried out with protocols equivalent to the ones described for expression of *Ct*UGGT(42). The immobilised metal affinity chromatography (IMAC) purification step was carried out in an equivalent manner to the same IMAC step for the purification of *Ct*UGGT as previously described(42), except that a 20 Column Volumes gradient elution was carried out at a flow rate of 1 ml/min increasing from 0% to 100% elution buffer.

Size Exclusion Chromatography (SEC): the IMAC step eluate was pooled and concentrated to 0.5 mL using a 100kDa spin concentrator. The sample was then loaded on a 0.5 mL loop and applied to a 10/300 Sephadex 200 column running at 1 mL/min. The SEC buffer was 20 mM MES pH 6.5, 50 mM NaCl, 1 mM CaCl_2_, 1 mM UDP. The latter buffer was arrived at by Differential Scanning Fluorimetry (DSF): the stability of UGGT1 is greatly increased through the addition of CaCl_2_, with an increase in melting temperature Tm of 3.0 °C and addition of UDP, with an increase in Tm of 1.1 °C. The DSF experiment also showed a clear preference for lower salt concentrations and a slightly more acidic pH.

### Ct UGGT_GT24_ cloning, protein expression and purification

The DNA encoding C-terminally His-tagged *Ct*UGGT_GT24_ (residues 1187-1473) was successfully amplified by PCR starting from the *Ct*UGGT-pHLSsec vector(42) using primers *Ct*UGGT_GT24__Fwd: ggttgcgtagctgaaaccggtGAGGCAACCAAGTCCGTG and *Ct*UGGT_GT24__Rev: gatggtggtgcttggtaccTTCCCTCACTCTCCTCGC.

The amplified insert was identified by agarose gel electrophoresis 900 bp and purified from the gel.

Following purification of the PCR products, the *Ct*UGGT_GT24_ insert was assembled into the AgeI/KpnI linearised pHLSec vector via ligation independent cloning (*aka* Gibson assembly). After transformation and plating, *E*.*coli* colonies containing the desired construct were identified by colony PCR through identification by agarose gel electrophoresis of the correct size of 900. *Ct*UGGT-pHLsec DNA plasmid purification from the correctly identified colonies was carried out via DNA miniprep and the resulting plasmid DNA sent for sequencing for confirmation of the desired DNA construct.

The maxiprepped *Ct*UGGT-pHLsec DNA plasmid was transfected into HEK293F cells following the protocol used for *Ct*UGGT(42). Purification was achieved by IMAC on an Åkta FPLC system, followed by gel filtration chromatography, after which the proteins were identified by SDS-PAGE. The final buffer was 20 mM HEPES pH 7.4, 50 mM NaCl, 1 mM CaCl_2_.

### Crystal growth

Crystals were grown at 18 °C in sitting drops by the vapour diffusion method, set up with a Mosquito liquid handling robot (TTP Labtech). Crystallisation drops had an initial volume of 200 nL. The volume ratio of protein to precipitant was either 1:1 or 2:1.

### CtUGGT_GT24_ crystallisation

A crystal of *Ct*UGGT_GT24_ grew in one week in a 1:1 mixture of *Ct*UGGT at 6 mg/mL and Morpheus screen condition 1-1 composed of 0.06M Divalents, 0.1 M Buffer System 1 pH 6.5, 30% v/v Precipitant Mix 1(62, 63).

### CtUGGT_GT24_:U2F crystallisation

U2F was synthesised as described(64). A crystal of *Ct*UGGT_GT24_:U2F grew in one week in a 1:1 mixture of *Ct*UGGT at 12 mg/mL, 2 mM CaCl_2_, 1.25 mM U2F and Morpheus screen condition 2-17 composed of 0.12 M Monosaccharides, 0.1 M Buffer System 2 pH 7.5, 30% v/v Precipitant Mix 1(62, 63).

### 5M-8OH-QCtUGGT_GT24_ crystallisation

A crystal of ^5M-8OH-Q^*Ct*UGGT_GT24_ grew in one week in a 1:1 mixture of *Ct*UGGT at 6.5 mg/mL, 10 mM 5M-8OH-Q in DMSO and Morpheus screen condition 1-1 composed of 0.06M Divalents, 0.1 M Buffer System 1 pH 6.5, 30% v/v Precipitant Mix 1(62, 63).

### *Ct*UGGT_GT24_ X-ray data collection, processing, and model refinement

X-ray data collection beamlines are listed in Table S1. Data processing was carried out in autoPROC(65). The model refinement and ligand fitting were carried out with BUSTER(66, 67) and Coot(68, 69). The highest resolution structure (UDP-Glc conformation 1) was used to provide external restraints for the refinement of the other structures; the P1 structure (5M-8OH-Q) was refined with additional automated NCS restraints (70).

### Measurement of the 5M-8OH-Q: human UGGT1 *K*_*d*_ dissociation constant by saturation transfer difference (STD) NMR *in vitro*

A 1 *µ*M solution of human UGGT1 was incubated with 5M-8OH-Q in PBS prepared in D_2_O and STD measured by NMR. The concentration dependence of the STD amplification factor (STD_*amp*_) was measured for each of the 5 aromatic hydrogen atoms in 5M-8OH-Q. The fit to the five sets of data pertaining to each of the 5M-8OH-Q H nuclei was compatible with a single value of *K*_*d*_ (as expected, since all five H nuclei are part of the same ligand molecule). The *K*_*d*_ of best fit was 613.1 *µ*M. Epitope mapping of the STD data is in good agreement with the crystal structure, with protons in the A and E environments having the largest STD_*amp*_ values, indicating they are in closest proximity to the protein when bound.

### *In cellula* UGGT-mediated glucosylation assays

The *in cellula* UGGT-mediated glucosylation assays were carried out as previously described(35), in presence of increasing amounts of 5M-8OH-Q. Briefly, HEK293-6E cells were plated and grown for 24 hr before replacing with fresh media containing the drug. After a 5 hr incubation time, the media was collected and the adhered cells were removed from the plate with lysis buffer. The media fraction was gently spun down (250xg for 5 min) to collect the dissociated cells and combined with the cells scraped off the plate. The combined samples were then shaken for 10 min at 4 °C before being spun at 14,000xg for 10 min at 4 °C prior to analyzing the soluble fraction.

### *In silico* docking of the Man_9_GlcNAc_2_ glycan on the surface of the *Ct*UGGT_GT24_ domain

To build an *in silico* model of the Man_9_GlcNAc_2_ Glc_1_ glycan bound to the *Ct*UGGT catalytic domain, we used a hierarchical approach that combined biased docking and Molecular Dynamics (MD). This protocol overcomes the limitation of conventional docking methods in dealing with polysaccharide ligands larger than five units (71). As a rule, carbohydrate ligands bind to proteins in a conformation close to one of the gas-phase energy minima. The latter mainly depend on the values of the dihedral angles of each glycosidic bond(72). Although each of these can only assume a few possible conformations, their number is such that docking algorithms cannot handle so many of degrees of freedom (73). For this reason, we first generated many Man_9_GlcNAc_2_ conformations using the `GLYCAM-web` server at (www.glycam.org); then, each structure was energy-minimised using MD in explicit solvents with the carbohydrate specific `GLYCAM06` force field (74). The results were clustered using only the furanose ring with a 1.4 A of tolerance (BlancoCapurro et al. 2019) and representative Man_9_GlcNAc_2_ conformations for each cluster were selected for the following steps:

1. we first aligned the acceptor Man residue of the Man_9_GlcNAc_2_ N-linked glycan (i.e. the terminal Man residue of its A-branch, Man “G” (see Figure 4B) such that its C1 atom pointed towards the O3 atom of the UDP-Glc molecule in our ^U2F^*Ct*UGGT_GT24_ structure and the structure of *Td*UGGT_*GT* 24_in complex with UDP-Glc (PDB ID 5H18, (26)). This assumes that this Man “G” residue docks in the active site such that upon Glc transfer, a *β*(1-3) linkage will form;
2. then, using that Man “G” residue orientation as a constraint, we performed multiple docking simulations of the Man_9_GlcNAc_2_ ligand, using the `AutoDock-Bias` protocol ((75) modified as described in (71);
3. the results were clustered and the three best ranking poses selected for further refinement using MD simulations. Starting from each complex, Molecular Dynamics was used to relax the Man_9_GlcNAc_2_ structure onto the *Ct*UGGT_GT24_ domain, using the protocol described in (76);
4. since the final pose for each of the three MD refinements was almost identical (RMSD < 2Å), we performed a final single-point energy calculation with `AutoDock4` (77)(Morris et al. 2009) to select the best complex.

## ACKNOWLEDGMENTS

We thank the members of N.Z.’s laboratory for assistance in the lab. Edward Lowe, Patrick Collins, Alice Douangamath, Jose Brandao-Neto and the staff at beamline I04-1, at the Diamond Light Source, Harwell, England, UK assisted with crystal growing, soaking, fishing and X-ray data collection. Jo Nettleship at the Oxford Protein Production Facility assisted in the cloning of UGGT1. Christina Redfield assisted with the NMR measurements. The work was funded by the Glycobiology Endowment and by a University of Oxford Confidence in Concept Scheme, grant reference MRC–MC_PC_16056 (to N.Z.). P.R. was the recipient of an LISCB Wellcome Trust ISSF award, grant reference 204801/Z/16/Z and a Wellcome Trust Seed Award in Science, grant reference 214090/Z/18/Z. This work was also supported by the National Institutes of Health (GM086874 to D.N.H.) and a Chemistry-Biology Interface program training grant (T32 GM139789 to K.P.G)). R.I. was the recipient of Sardinian Regional Government Erasmus and PhD scholarships. N.Z. is a Fellow of Merton College, Oxford.

## Footnotes

* During the synthesis of the N-linked glycan precursor, ALG6 appends the first glucose to the Man_9_GlcNAc_2_ carbohydrate - before it is built into a triglucosylated form by other ALG enzymes at the ER membrane. The Glc_3_Man_9_GlcNAc_2_ glycan is then appended to newly synthesized proteins and trimmed to a monoglucosylated state by glucosidases I and II, providing the ligand necessary for binding to the lectin chaperones calnexin and calreticulin (37). During glycan maturation in wild type cells, ER lectins binding can occur after the glycan is trimmed from a Glc_3_Man_9_GlcNAc_2_ to a GlcMan_9_GlcNAc_2_ form, or through glucosylation of a Man_9_GlcNAc_2_ carbohydrate by the UGGTs (36).

