## Supplemental Figures and Tables for "A quinolin-8-ol sub-millimolar inhibitor of UGGT, the ER glycoprotein folding quality control checkpoint"

PNAS
www.pnas.org

### Supplementary Information for

#### A quinolin-8-ol sub-millimolar inhibitor of UGGT, the ER glycoprotein folding quality control checkpoint

**Kevin P. Guay, Roberta Ibba, JL Kiappes, Maria De Benedictis, Ilaria Zeni, James D. Le Cornu, Mario Hensen, Anu V. Chandran, Anastassia L. Kantsadi, Alessandro T. Caputo, Juan I. Blanco Capurro, Yusupha Bayo, Johan C. Hill, Kieran Hudson, Andrea Lia, Snežana Vasiljević, Carlos P. Modenutti, Stephen G. Withers, Marcelo Martí, Emiliano Biasini, Angelo Santino, Daniel N. Hebert, Nicole Zitzmann, Pietro Roversi**

**Pietro Roversi**
****

##### This PDF file includes:

Figs. S1 to S7
Tables S1 to S3
SI References

#### Supplementary Figures

##### Methods.

**Viability assay for treated HEK293-6E cells.** . The viability of cells after drug treatment was determined using a LUNA II™ Automated Cell Counter. Briefly, untreated and treated cells were incubated with the drug 5 hr. After incubation cells were collected and washed twice with PBS and resuspended in 1 mL of culture media. Cells were mixed with trypan blue (50:50 mix) and viability was measured.

**Treatment of *Arabidopsis thaliana* with 5M-8OH-Q.** Seeds of *Arabidopsis thaliana* WT, ecotype Columbia were sown on plates containing half-strength Murashige and Skoog (MS) agar medium. Seedlings were then transferred to small plates containing liquid MS medium and plants grown in these conditions for 10 days before treatment with 2mM of 5M-8OH-Q for 2, 4, 6 and 12h. N-glycosylated proteins were detected by digestion of total protein extracts with Endoglicosidase H (Sigma-Aldrich), followed by blot using 2 $\mu$g/mL of peroxidase-conjugated concanavalin A (Sigma-Aldrich) following the manufacturer's recommendations.

**Propidium iodide dye staining of *Arabidopsis thaliana* for 5M-8OH-Q toxicity assay.** WT seedlings treated with 5M-8OH-Q for indicated times were stained with 10 $\mu$g/mL propidium iodide (PI; Sigma-Aldrich) solution for 10 minutes and then washed with distilled water. Root tip samples were observed with a confocal laser-microscope (LSM Pascal, Zeiss, ZEN Software) with filter set (excitation 488 nm, emission $\geq$ 543 nm). Red fluorescence of cell wall indicates alive cells, while nuclear staining indicates damaged cells.

**Treatment of HEK293 cells with 5M-8OH-Q and anti-PrP immunoblotting of protein fractions.** For PrP detection in the Western blot assay, cells were seeded in a 24-wells plate at 100,000 cells/well. After 24 hours, the whole medium was replaced and cells were treated with 1 mM 5M-8OH-Q (using a 300 mM stock solution in DMSO). Cells were harvested at different time points: 4, 8 and 24 hours after treatment. Cells harvested at each time point were lysed with 30 $\mu$L of lysis buffer (Tris 20 mM, NaCl 150 mM, 0.5% Triton X-100, 0.5% NP40) containing phosphatase and protease inhibitors.

Protein quantification was performed using the BCA Assay Kit (Pierce), as per manufacturer's instructions. For each sample, an aliquot corresponding to 21 $\mu$g of total proteins was diluted in 4X Laemmli Buffer (Bio-Rad) containing 100 mM dithiothreitol (DTT). Samples were denatured by heating at 95°C for 8 minutes, loaded on a Bio-Rad precast protein gradient SDS-PAGE gel and electrophoresis run at constant voltage 135 V. Proteins were then transferred to a polyvinylidene fluoride (PVDF) membrane and blotted with an anti-PrP antibody (D18) diluted in BSA 3% (w/v) in T-TBS (Tris-Buffered Saline, 0,1% Tween) overnight at 4°C. Signals were revealed with a horseradish conjugated goat anti-human IgG diluted 1:5000 for 1 h at room temperature and acquired with a ChemiDoc XRS Touch Imaging System (Bio-Rad). The final quantification of proteins detected by the primary antibody was obtained by densitometric analysis of the western blots, normalizing each signal on the corresponding total protein lane (obtained by the enhanced tryptophan fluorescence technology of stain-free gels, BioRad).

PnGase F treatment: The PnGase F treatment kit (New England Biolabs, Beverly, MA, USA) was used according to manufacturer's instructions. Briefly, denaturating buffer was added to a 30 $\mu$g aliquot of protein lysates diluted in 18 $\mu$L of lysis buffer (Tris 20 mM, NaCl 150 mM, 0.5% Triton X-100, 0.5% NP40). Samples were then denatured for 10 minutes at 95 °C. After 2 minutes on ice, GlycoBuffer 2 (New England Biolabs) and NP40 1% were added. Each sample was split in two aliquots, one incubated with PnGase F(125 U) and the other with an equivalent volume of dd$H_2O$, and incubated for 1 hour at 37°C. The samples were then analyzed by western blot assay as described above.

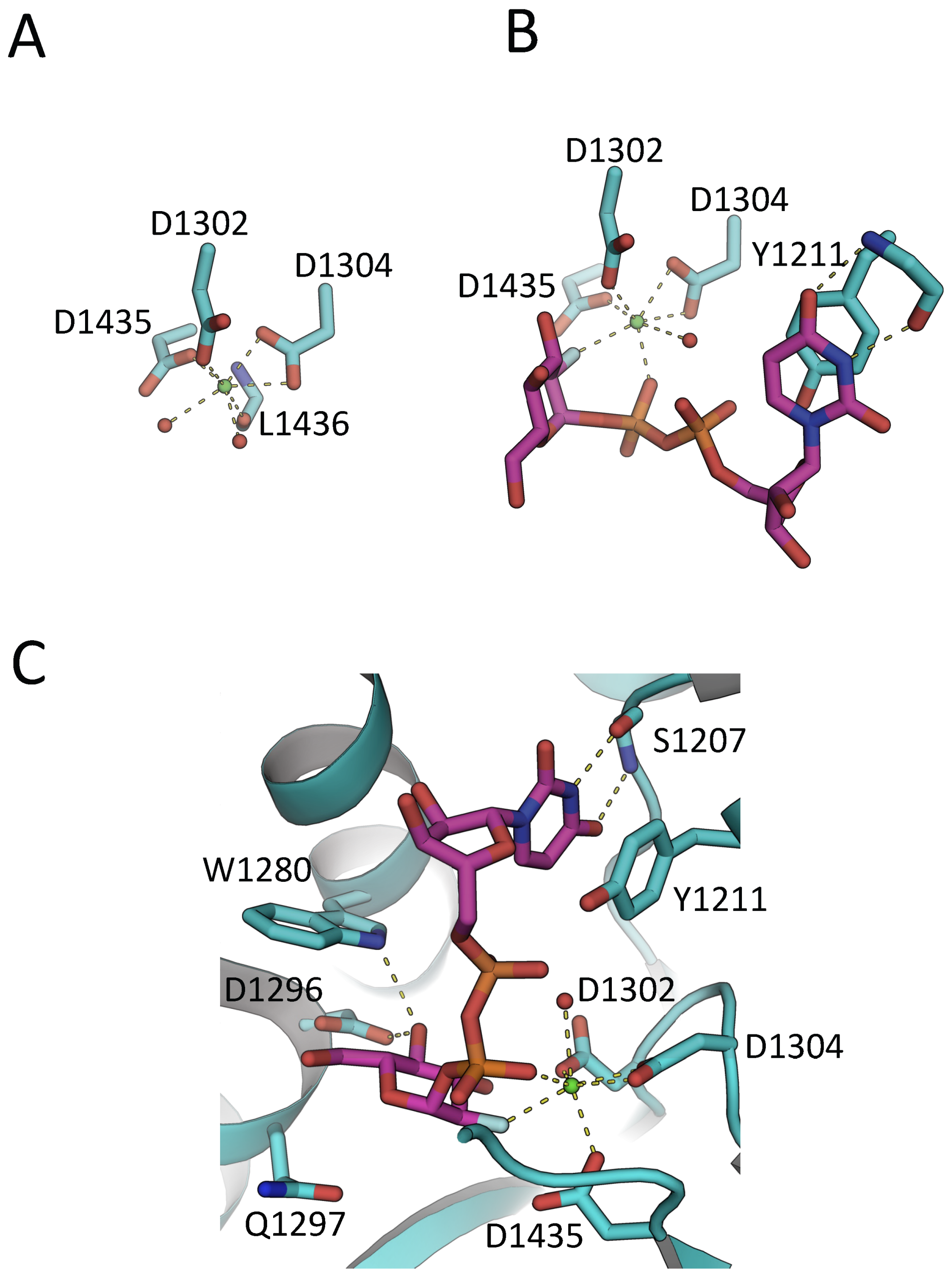


**Fig. S1. The active sites of *Ct*UGGT$_{GT24}$ and *Ct*UGGT$_{GT24}$$^{U2F}$.** Protein atoms in sticks representation; C cyan (but UDP-Glc(UDP) C magenta and 5M-8OH-Q C atoms yellow), O red, N blue, P orange, F light green. H-bonds and $Ca^{2+}$-coordination bonds are in yellow dashed lines. The $Ca^{2+}$ ion is a green sphere and its coordinating water molecules are red spheres. The side chains of residues D1302, D1304 and D1435 coordinate the $Ca^{2+}$. **A**: apo *Ct*UGGT$_{GT24}$ (PDB ID 7ZKC). The octahedral coordination sphere of the $Ca^{2+}$ ion is completed by two water molecules and the main chain of L1436. **B**: *Ct*UGGT$_{GT24}$$^{U2F}$(PDB ID 7ZLU). L1436 moves away from the $Ca^{2+}$ ion, and two coordination sites are taken up by the U2F $\beta$ phosphate and the F atom at position 2' of the Glc ring. The uracyl O4 atom accepts a H-bond from the S1207 main chain NH. Only one $Ca^{2+}$-coordinating water molecule remains. On the right hand side, the residues coordinating the uracyl ring: the side chain of *Ct*UGGT$_{GT24}$ Y1211 and the main chain of S1207. **C**: the UGGT active site selects UDP-Glc over UDP-Gal (1–3): in UDP-Glc the glucose O4 atom forms hydrogen bonds to the side chains of conserved W1280 and D1396 (Figure S1-C), but these interactions would be lost in UDP-Gal (because of the difference in stereochemistry between Glc and Gal in position 4).

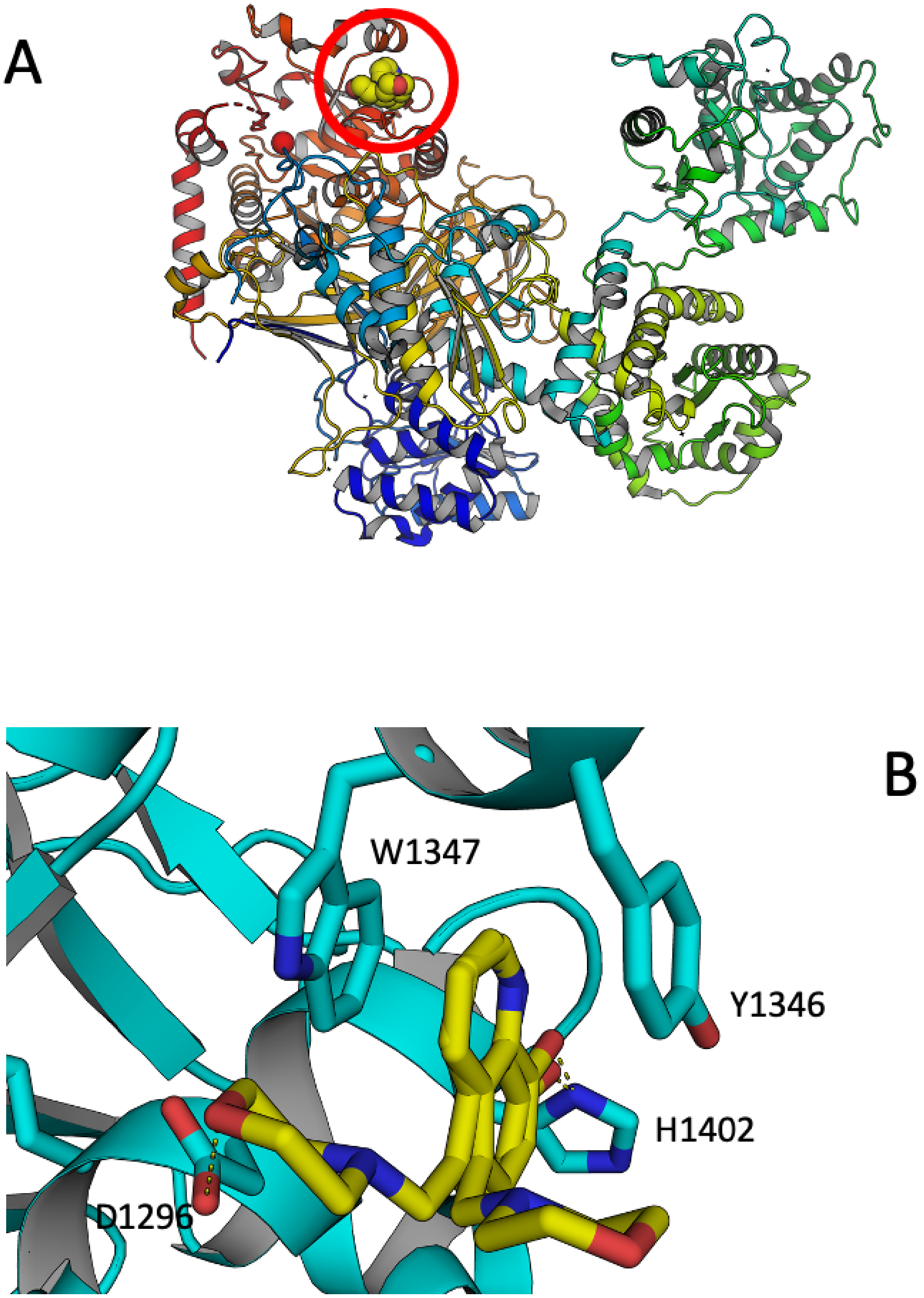


**Fig. S2.** **A**: the structure of *Ct*UGGT$_{GT24}^{5M\text{-}8OH\text{-}Q}$(PDB ID 7ZLL) in complex with 5M-8OH-Q, superposed onto the structure of *Ct*UGGT (PDB ID 5MZO) in order to illustrate the binding site of the inhibitor in the context of the whole structure. The protein is in cartoon representation coloured blue-to-red from N- to C-terminus; the 5M-8OH-Q is inside a red circle, in spheres representations (C atoms in yellow). **B**: Zoom onto the *Ct*UGGT $^{1346}$YW$^{1347}$ clamp (C atoms in green) binding 5M-8OH-Q (C atoms in cyan) Representative distances to interacting residues are in dashed lines. Only two of the many morpholine ring placements are shown. PDB ID: 7ZLL.

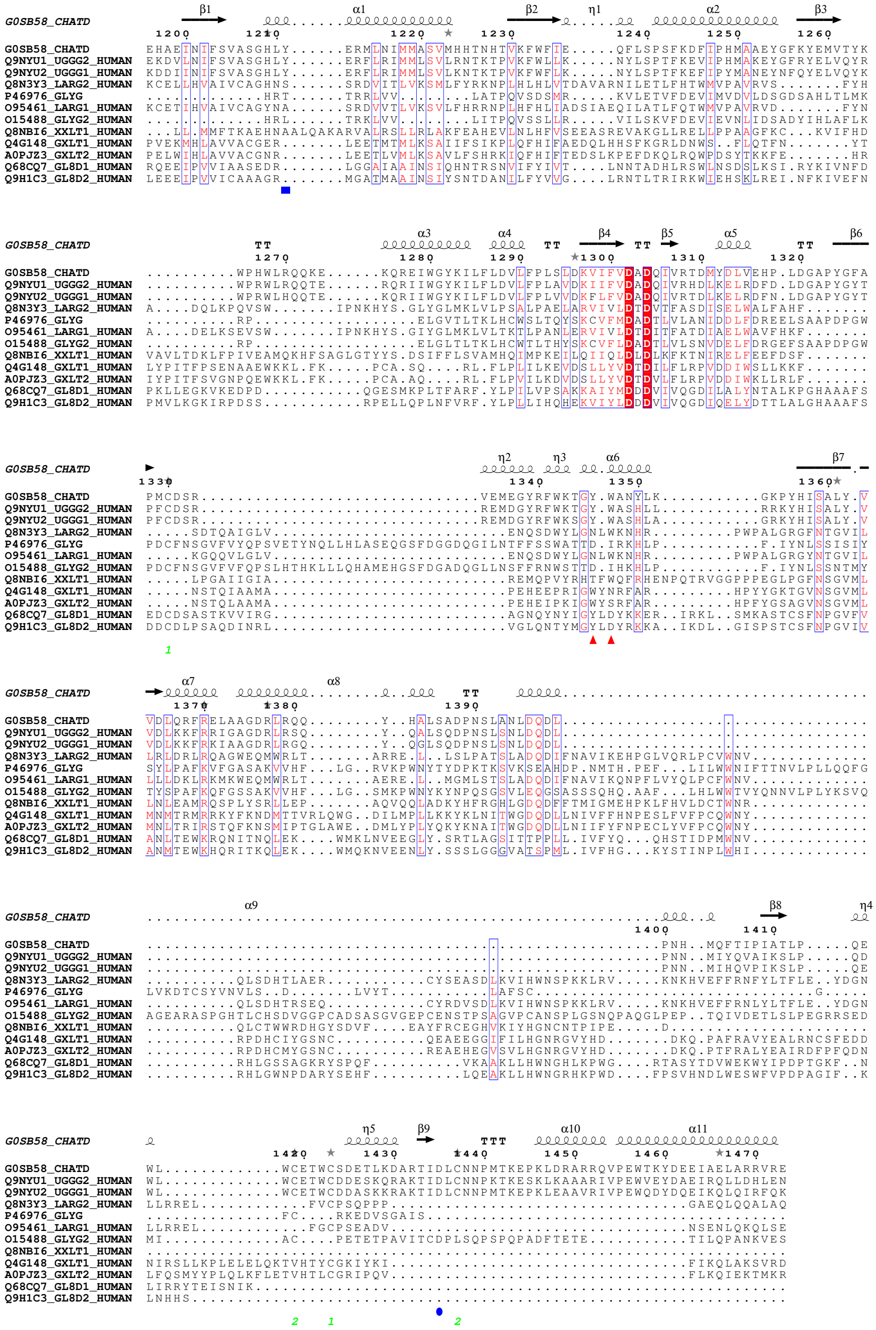

**Fig. S3. Alignment of GT-24 and GT-8 domains in human proteins**. Uniprot codes and proteins: Q8N3Y3, Xylosyl- and glucuronyltransferase LARGE2; P46976, Glycogenin 1; O95461, Xylosyl- and glucuronyl-transferase LARGE1; O15488, Glycogenin 2; Q8NBI6, Xyloside xylosyltransferase 1; Q4G148, Glucoside xylosyltransferase 1; A0PJZ3, Glucoside xylosyltransferase 2; Q68CQ7, Glycosyltransferase 8 domain-containing protein 1; Q9H1C3, Glycosyltransferase 8 domain-containing protein 2. The *Ct*UGGT D1302 and D1304 residues coordinating $Ca^{2+}$ ion are completely conserved across these sequences. Red triangles mark the *Ct*UGGT $^{1346}$WY$^{1347}$ clamp. A blue square marks the position of *Ct*UGGT Y1211 (coordinating the U2F uracyl ring). A blue oval marks the position of *Ct*UGGT D1435 (coordinating the $Ca^{2+}$ ion).

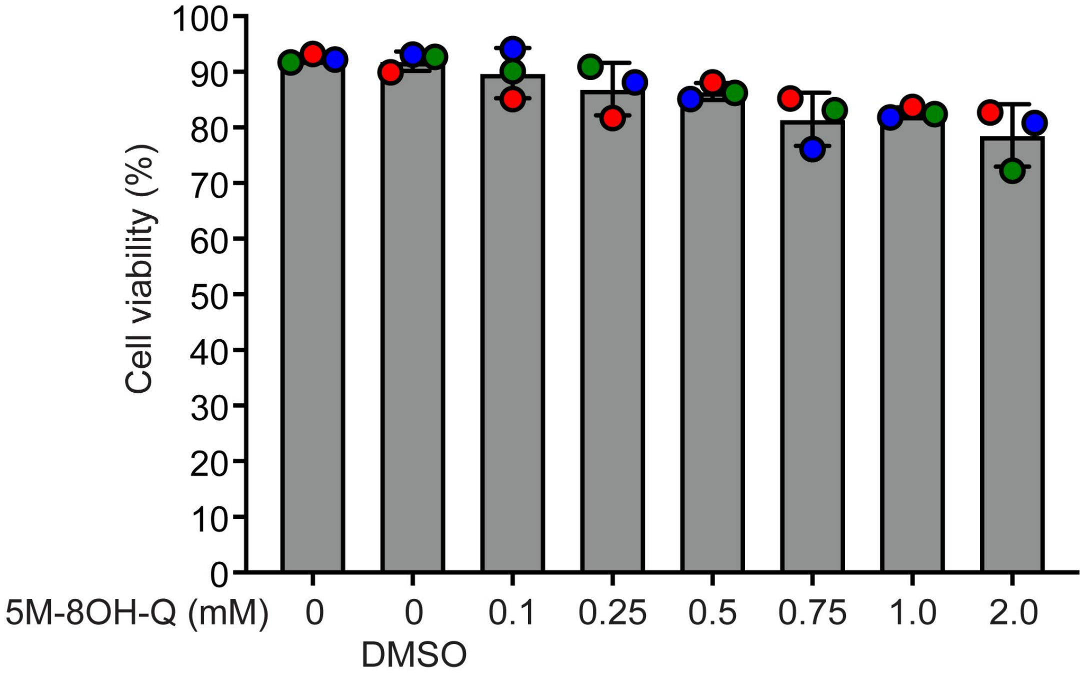


**Fig. S4. Viability of *ALG6*$^{-/-}$ HEK293-6E cells after 5 hr 5M-8OH-Q treatment.** Prior to lysis, cells were dissociated and washed twice with PBS before being resuspending in 1 mL of PBS. Resuspended cells were diluted 1:1 with trypan blue and counted using a LUNA II$^{TM}$ Automated Cell Counter. Error bars represent the standard deviation of three independent biological replicates.

#### 1. Supplementary Tables

**Table S1. X-ray data collection parameters and data processing statistics for *Ct*UGGT$_{GT24}$ crystal structures.**

| Structure | *Ct*UGGT$_{GT24}$ | *Ct*UGGT$_{GT24}$$^{U2F}$ | *Ct*UGGT$_{GT24}$$^{5M-8OH-Q}$ |
|---|---|---|---|
| PDB ID | 7ZKC | 7ZLU | 7ZLL |
| Beamline | I03@DLS | I04@DLS | I04@DLS |
| Wavelength $\lambda$ (mm, Å) | 0.97960 | 0.97956 | 0.97950 |
| Transmission % | 100 | 100 | 100 |
| Number of images | 1,800 | 1,800 | 3,600 |
| Oscillation range (°) | 0.1 | 0.1 | 0.1 |
| Exposure time (s) | 0.02 | 0.017 | 0.1 |
| Space Group (Z) | H3 (6) | P1 (3) | H3 (6) |
| Cell edges: a,b,c (Å) | a=b=118.82,c=62.12 | a=68.16,b=72.53, c=72.39 | a=b=118.858, c=68.551 |
| Cell angles $\alpha, \beta, \gamma$ (°) | $\alpha$=$\beta$=90, $\gamma$=120 | $\alpha$=110.72, $\beta$=108.33, $\gamma$=108.27 | $\alpha$=$\beta$=90, $\gamma$=120 |
| Resolution Range (Å) | 59.41-1.77 (1.86-1.77) | 41.16 - 2.05 (2.32-2.05) | 41.16-1.65 (1.74-1.65) |
| $R_{merge}$ | 0.14 (2.01) | 0.05 (0.42) | 0.05 (1.60) |
| $R_{meas}$ | 0.15 (2.25) | 0.08 (0.60) | 0.05 (1.67) |
| Observations | 322,614 (25,624) | 68,743 (3,486) | 404,079 (21,046) |
| Unique observations | 35,287 (5,115) | 39,166 (1,958) | 38,495 (1,925) |
| Average I/$\sigma$(I) | 7.8 (0.7) | 8.4 (1.6) | 20.8 (1.3) |
| Completeness % | 99.7 (99.2) | 81.1 (45.9) | 88.4 (28.7) |
| Multiplicity | 9.1 (5.0) | 1.8 (1.8) | 10.5 (10.9) |
| $CC_{1/2}$ | 0.99 (0.35) | 0.998 (0.646) | 1.000 (0.581) |

Each structure was determined using a single crystal. Values in parentheses refer to the highest resolution shell.

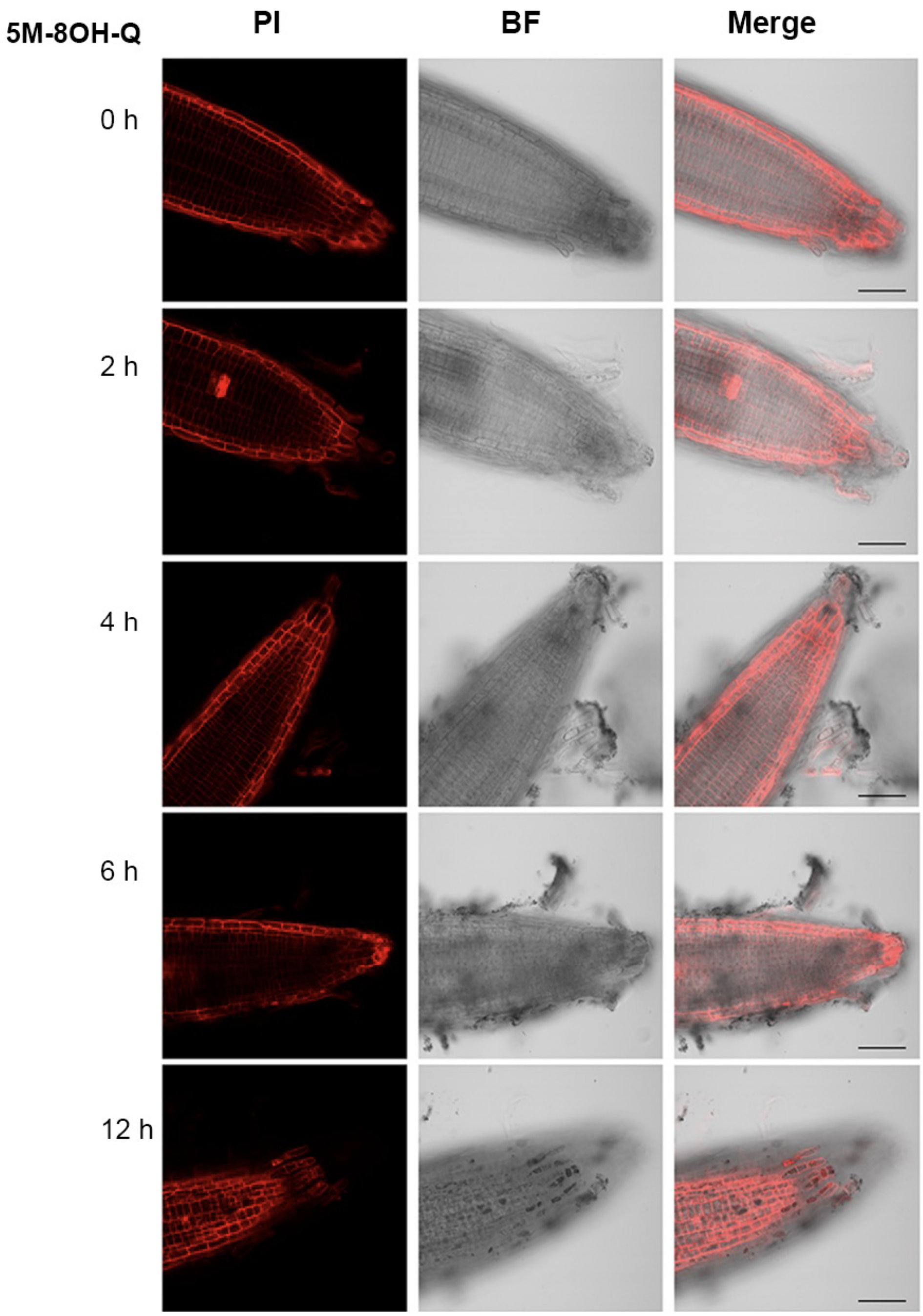


**Fig. S5. Effects of 5M-8OH-Qtreatment on cell viability of *A. thaliana* WT seedlings**. 10-days-old *A. thaliana* WT seedlings were treated for 2, 4, 6 and 12 hours with 2 mM inhibitor and root tips were stained with propidium iodide (PI). Confocal images shown the red fluorescence of PI, the bright field (BF) and merge. Scale bars, 50 $\mu$m. Red fluorescence of cell wall indicates alive cells, while nuclear staining indicates damaged cells

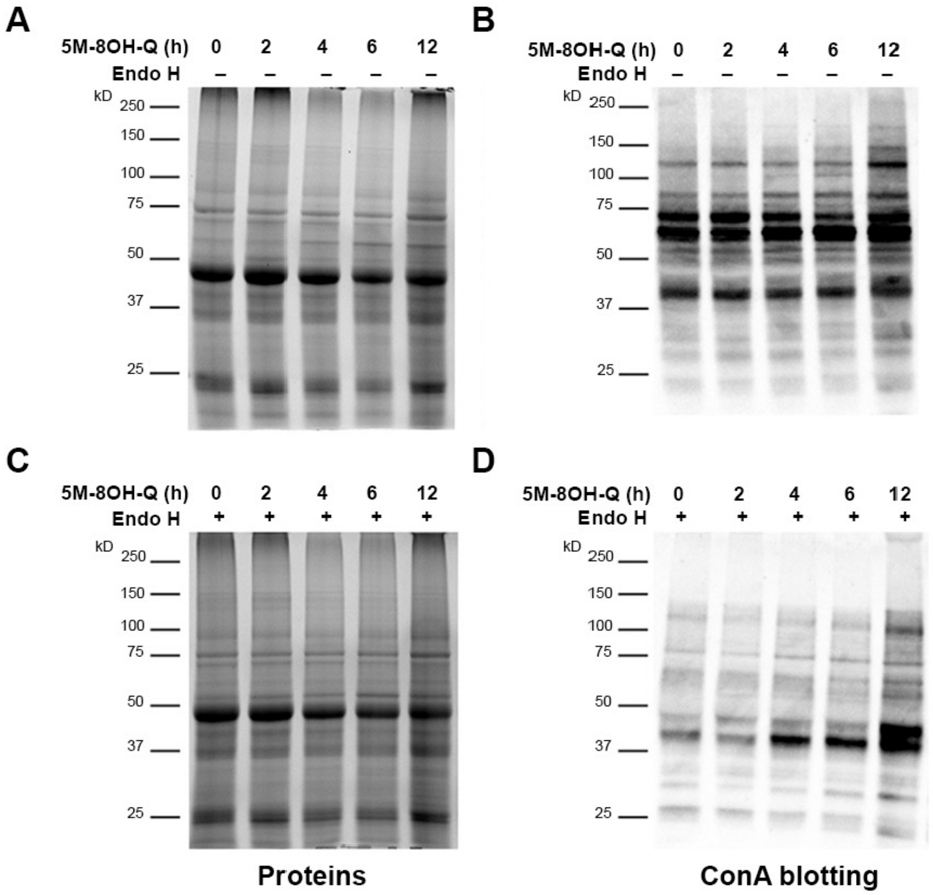


**Fig. S6. 5M-8OH-Q affects N-glycosylation of proteins in planta.** Seedlings of *At* WT treated with 5M-8OH-Q were used to extract total protein, followed by deglycosylation treatment with Endoglycosidase H (Endo H) , then separated by SDS-PAGE (**A,C**), blotted and treated with concanavalin A (ConA) for high mannose N-glycan detection (**B,D**).

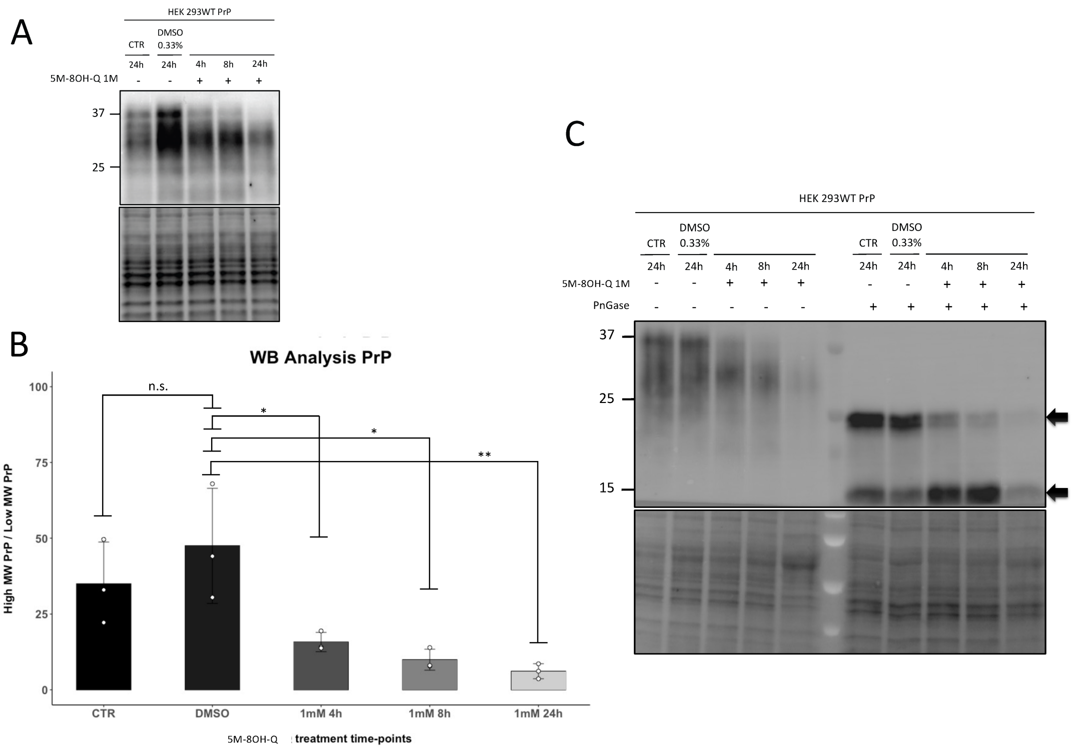


**Fig. S7. Inhibition of UGGT increases the alpha-cleavage of PrP.** Prion diseases are fatal neurodegenerative disorders affecting humans and several animal species. They are caused by the aggregation of a misfolded isoform (called PrPSc) of the major prion protein (PrP), a cell-surface glycoprotein of uncertain function. PrPC biogenesis follows a trafficking pathway typical of glycosylphosphatidylinositol (GPI)- anchored polypeptides. The protein is synthesized directly in the lumen of the endoplasmic reticulum (ER), where it folds and receives post-translational processing of the primary structure. These include the removal of signal peptide at the N-terminus and the addition of a GPI anchor as well as two N-linked glycans (at Asn-181 and Asn-197) at the C-terminus. Most PrP is found on the cell-surface where it is localized to lipid rafts, although a fraction is endocytosed via clathrin-coated pits. Some of the protein is proteolytically cleaved in the endosomal recycling pathway by members of the ADAM metalloproteases near residue 111 to generate N- and C-terminal fragments called N1 and C1, respectively (4). In order to evaluate the effect of 5M-8OH-Q on the homeostasis of PrP, we employed HEK293, which express very low levels of the protein endogenously, stably transfected with wild type (WT) PrP under the control of a CMV promoter. Cells exposed to 5M-8OH-Q (1 mM), vehicle (DMSO, volume equivalent) or untreated (CTR) were analyzed at different time points by western blotting. We observed a time-dependent reduction of mature, full-length PrP after treatment with 5M-8OH-Q (Panels **A**,**B**). In order to assess whether such a decrease was reflecting an altered glycosylation or an increase alpha cleavage, we repeated the experiment by adding a step of de-glycosilation before the Western blotting (Panel **C**). We observed that 5M-8OH-Q induces a time-dependent decrease of full-length PrP and a parallel increase of the C1 fragment. Thus, 5M-8OH-Q strongly increases the alpha cleavage of PrP. Since UGGT regulates several proteins transiting through the ER, our data suggest that a client of UGGT could directly or indirectly influence the function of ADAM metalloproteases involved in PrP's alpha cleavage. **A**: HEK293 cells stably expressing WT PrP were treated with 5M-8OH-Q at a final concentration of 1 mM for three different time points (indicated). The total proteins obtained from each cell lysate were loaded onto an SDS-Page gel for the western blotting analysis. The PrP signal was revealed with an anti-PrP antibody (D18). **B**: Graph shows the ratio between the densitometric quantification of the higher molecular weight (MW) and the low MW PrP bands. Each signal was normalized on the corresponding total protein lane (detected by UV) and expressed as a percentage value. Three independent replicates were performed (each value is represented as a white dot), and the mean was calculated and reported in the bar plot. The error bars reflect the standard deviation. Student t-test was performed for statistical analysis: n.s.:non-significant; *: $P \leq 0.05$, **: $P \leq 0.02$. **C**: HEK293 cells stably expressing WT PrP were treated with 5M-8OH-Q at a final concentration of 1 mM for three different time points (indicated). The total proteins obtained from each cell lysate were split in two aliquots, one incubated with PnGase F (125 U) and the other with an equivalent volume of $ddH_2O$. The PrP signal was revealed with an anti-PrP antibody (D18) recognizing an epitope in the C-terminus of the protein downstream of the alpha cleavage, thus allowing the visualization of full-length PrP and the C1 fragment. Arrows indicate the two different PrP forms.

**Table S2. Refinement statistics for *Ct*UGGT$_{GT24}$ crystal structures. All structures contain a $Ca^+$ ion coming from the protein solution.**

| Structure | *Ct*UGGT$_{GT24}$ | *Ct*UGGT$_{GT24}$$^{U2F}$ | *Ct*UGGT$_{GT24}$$^{5M\text{-}8OH\text{-}Q}$ |
|---|---|---|---|
| PDB ID | 7ZKC | 7ZLU | 7ZLL |
| Ligand | - | U2F | 5M-8OH-Q |
| Wavelength $\lambda$ (Å) | 0.97960 | 0.97956 | 0.97950 |
| Space Group (Z) | H3 (6) | P1 (3) | H3 (6) |
| Resolution Range (Å) | 59.41-1.77 (1.86-1.77) | 41.16 - 2.05 (2.32-2.05) | 59.27-2.53 (2.67-2.53) |
| $R_{work}$,$R_{free}$ | 0.205, 0.226 (0.317,0.331) | 0.226, 0.270 (0.322, 0.445) | 0.209, 0.235 (0.310, 0.301) |
| Protein atoms ($<$B factor$>$, Å$^2$) | 2,432 (43.64) | 7,137 (36.44) | 2,448 (38.34) |
| Water molecules ($<$B factor$>$, Å$^2$) | 227 (48.48) | 289 (34.65) | 226 (48.00) |
| Ligands ($<$B factor$>$, Å$^2$) | $Ca^{2+}$ (30.34) | 3 $\times$ ($Ca^{2+}$, U2F) (42.62) | $Ca^{2+}$, 5M-8OH-Q (49.56) |
| $rmsd_{bonds}$ (Å), $rmsd_{angles}$ ($^\circ$) | 0.008, 0.90 | 0.008, 0.94 | 0.008, 0.93 |
| Number of Ramachandran favoured (% ) | 268 (100.0) | 824 (98.0) | 287 (99.0) |
| Number of Ramachandran allowed (% ) | 1 (0.0) | 15 (2.0) | 3 (1.0) |
| Number of Ramachandran outliers (% ) | 0 (0.0) | 0 (0.0) | 0 (0.0) |

Each structure was determined using a single crystal. Values in parentheses refer to the highest resolution shell.

**Table S3. Mammalian protein structures with 8-OH-quinoline ligands.**

| PDB ID (Ref.) | Protein | Uniprot ID | 8-OH-Q ligand (PDB name) | Binding site |
|---|---|---|---|---|
| 3KCY (5) | Hypoxia-inducible factor 1-alpha inhibitor (HIF1AN) | Q9NWT6 | " | $Fe^{++}$ |
| 4BIO, 3OD4 (6) | " | " | 8-hydroxyquinoline-5-carboxylic acid (8XQ) | $Zn^{++}$ or $Fe^{++}$ |
| 2XXZ | Lysine-specific demethylase 6B (KDM6B) | O15054 | " | $Ni^{++}$ instead of physiological $Fe^{++}$ |
| 3NJY (7) | Lysine-specific demethylase 4A (KDM4A) | O75164 | " | " |
| 6FUK (8) | Lysine-specific demethylase 6A (UTX) | O15550 | " | $Fe^{++}$ |
| 4IE4 (9) | Fat mass and obesity associated protein (FTO) | Q9C0B1 | " | $Zn^{++}$ |
| 4JHT (6) | *E. coli* Alpha-ketoglutarate-dependent dioxygenase (AlkB) | P05050 | " | $Mn^{++}$ |
| 6RBJ | Lysine-specific demethylase 3B (KDM3B) | H0Y946 | 5-(1 H-1,2,3,4-tetrazol-5-yl)quinolin-8-ol (JX8) | $Mn^{++}$ |
| 6RBI | Lysine-specific demethylase 5B (KDM5B) | Q9UGL1 | " | $Mn^{++}$ |
| 6AFR (10) | Bromodomain-containing protein 4 (BRD4) | O60885 | 5-[(4-fluoranylimidazol-1-yl)methyl]quinolin-8-ol (9E3) | quinolin-8-ol in hydrophobic pocket |
| 5Z5T (11) | " | " | 2-amino-4-(1H-imidazol-1-yl)quinolin-8-ol (96R) | " |
| 6LG7 | " | " | 2-azanyl-6-fluoranyl-4-imidazol-1-yl-quinolin-8-ol (ECF) | H-bond and hydrophobic interactions |
| 6LG8 | " | " | 2-azanyl-5-fluoranyl-4-imidazol-1-yl-quinolin-8-ol (ECR) | " |
| 6LG9 | " | " | 2-azanyl-7-bromanyl-4-imidazol-1-yl-quinolin-8-ol (ECU) | " |
| 4E26 (12) | Serine/threonine-protein kinase B-raf (BRAF) | P15056 | 5-chloro-7-[(R)-furan-2-yl(pyridin-2-ylamino)methyl]quinolin-8-ol (734) | quinolin-8-ol sandwiched between Trp and Phe rings |
| 5PA1 | *R. norvegicus* Catechol O-methyltransferase (Comt) | P15056 | 6-(4-fluorophenyl)quinolin-8-ol (7JS) | $Mg^{++}$ |
| 5PA7 | " | " | 6-(4-fluorophenyl)-8-oxidanyl-3 H-quinazolin-4-one (7JD) | $Mg^{++}$ |
| 6GY1 (13) | " | " | 7-fluoranyl-5-(4-methylphenyl)sulfonyl-quinolin-8-ol (FGQ) | $Mg^{++}$ |
| 3JSF (14) | Macrophage migration inhibitory factor (MIF) | P14174 | 7-(2-fluorobenzyl)quinolin-8-ol (XV1) | quinolin-8-ol sandwiched between Tyr and Phe rings |
| 3JSG (14) | " | " | 7-(pyridin-3-ylmethyl)quinolin-8-ol (0IN) | " |
| 3JTU (14) | " | " | 7-(pyridin-2-ylmethyl)quinolin-8-ol (ZIN) | " |

All proteins are human (*Homo sapiens*) unless otherwise specified.
